# Evaluating Evapotranspiration in a Commercial Greenhouse: A Comparative Study of Microclimatic Factors and Machine-Learning Algorithms

**DOI:** 10.1101/2024.01.11.575151

**Authors:** Nir Averbuch, Menachem Moshelion

## Abstract

The FAO-56 Penman-Monteith equation (FPME) is commonly used to calculate evapotranspiration and apply necessary irrigation, based on environmental data usually taken from a single measuring station. In this study, we hypothesized that the accuracy of the FPME is affected by microclimatic changes over time and space within the target area. Therefore, we tested the impact of numerous spatial and temporal environmental measurement points in a commercial greenhouse on the accuracy of the FPME, by comparing its evapotranspiration evaluation to the actual evaporation measured by dozens of weighing lysimeters throughout the year. Additionally, we harnessed the capabilities of machine-learning algorithms to utilize the extensive data acquired for predicting evapotranspiration. Our results revealed that the daily FPME exhibited a -22% to +22% discrepancy in accuracy, as compared to the lysimeters, with overestimation in the winter and underestimation in the summer. Interestingly, using more data points per day led to less accurate FPME evaluation. The conflict between an increased number of data points and a reduction in accuracy was explained by daily hysteresis. Machine-learning algorithms (Decision Tree, Random Forest, XGBoost and Neural Network) showed impressive accuracy in predicting evapotranspiration, when the model dataset contained temporal parameters (*R*^2^> 0.918). Furthermore, we demonstrated that spatial sampling had a stronger effect on the accuracy of predictions than the amount of the data collected. Specifically, when we used 10% of the original dataset (3.01e5 entries) with high consideration of spatial measurements, the best-performing models (Random Forest and XGBoost) were highly accurate (*R*^2^ = 0.913 and *R*^2^ = 0.935, respectively). The top three most influential features of all models were light, day and hour, underscoring the importance of the temporal dimension. This approach allowed us to explore the potential of leveraging advanced computational methods to improve the estimation of water loss under various environmental conditions.

## 1. Introduction

### 1.1. Background

Climate changes is expected to expose agriculture to a variety of challenges, from drought to flooding (Nunti et al., 2020). Moreover, population growth and increased standards of living necessitate increased food production (Van Dijk et al., 2021). The linear correlation between the amount of water applied to a crop and its yield (Liao et al., 2008) suggests that water is a limiting factor for crop growth and yield. In areas with insufficient or inconsistent precipitation, irrigation can play a key role. However, it can be difficult to determine the exact volume of irrigation water that should be applied. Therefore, farmers who have control over irrigation volumes tend to over-irrigate, in an effort to maximize crop yields. In addition to the financial cost involved, excessive irrigation also has its agronomic and environmental drawbacks, causing root diseases, soil salinization (Malash et al., 2008) and the contamination of ground water (Mahvi, 2005). Effective and precise irrigation is a challenging task due to spatial and temporal variation in meteorological and soil conditions (Ferentinos et al., 2017; Nebbali et al., 2012). Plants respond to these conditions instantaneously, by managing their water balance to optimize their productivity–survivability ratio (Markulj Kulundži et al., 2021). For example, adjustment of stomatal opening serves as a key physiological trait, optimizing the plant response by controlling the influx of CO2 and H20, to support photosynthesis and transpiration, respectively.

### 1.2. Indirect method to evaluate evapotranspiration

A better understanding of plant transpiration could help us to irrigate more efficiently. A few methods are used to evaluate evapotranspiration (*ET*_0_); the most cost-effective is the evaporation pan used as a reference for *ET*_0_ by applying an empirical crop factor (Haijun et al., 2015). The eddy-covariance method is also used to evaluate evapotranspiration, by frequently measuring the turbulence and gas exchange in fields and greenhouses. These figures are then translated into the evapotranspiration rate. However, the necessary measuring equipment is expensive and the practices are complex, with key assumptions that are challenging to apply in any agricultural area. The FAO-56 Penman-Monteith equation (FPME) was originally developed to evaluate the evaporation from well-irrigated, uniformly covering infinite grass (7 cm in height) on a flat surface. The FPME has been modified for use with different crop plants using empirical crop coefficient, as specific table was modified for each crop and his growth state. The FPME is considered the standard method for estimating evapotranspiration in greenhouses and agriculture fields (Ghiat et al., 2021). It involves meteorological parameters such as temperature, relative humidity (RH), vapor pressure deficit (VPD) and solar radiation. Data are usually collected by a single meteorological station serving a large area and daily or weekly spatial measurements are averaged, without any consideration of the short-term variability in microclimatic or physical characteristics (e.g., in the case of a greenhouse, the greenhouse walls, beams and unevenness in the moist mattresses). However, this variability in the growing environment of the individual plant has a significant impact on plant growth and evapotranspiration (Cui et al., 2016; Zheng et al., 2000). The averaging of the environmental variability may lead to inaccurate calculations of plant water use. This averaging can result in deviations in both the momentary and daily references to evapotranspiration. Evapotranspiration is also affected by the decoupling factor, which is the ratio between radiative and aerodynamic parameters (McNaughton & Jarvis et al., 1983). In a location in which those parameters are decoupled, solar radiation will account for most of the observed evaporation (Hadad et al., 2020; Haijun et al., 2015; Möller et al., 2004).

### 1.3. Direct method for evaluating evapotranspiration

Several methods exist for directly measuring evapotranspiration. Those methods aim to precisely determine water uptake and water availability for the plant. One such method involves the use of a dendrometer, which is a device that measures small changes in the diameter of a plant’s stem. However, this kind of apparatus cannot provide us with accurate figures for the amount of irrigation needed to compensate for transpiration losses (Clonch et al., 2021; Vilenski et al., 2019). Penetration methods are also used, such as thermal dissipation probes, which are placed inside the plant stem and measure water movement in the xylem by using passive heated electrical pulse. This method must be calibrated to provide good predictions of the absolute values of transpiration and inappropriate positioning of the probes (in terms of both radius or depth) will negatively affect the accuracy of their transpiration predictions (Granier, 1987; Strachan, 2016; Yin et al., 2014), as well as their predictions regarding the soil metric potential near the roots. This method is not considered accurate for determining the amount of irrigation needed. Since this method does not involve direct measurement of the plants, it is difficult to use this method to precisely evaluate the absolute amount of evapotranspiration. Weighing lysimeters (gravimetric systems) are widely regarded as accurate tools for monitoring changes in plants’ absolute transpiration, as described by (Halperin et al., 2017). Yet, weighing lysimeters are challenging to install in agricultural fields. In this study, we aimed to improve evapotranspiration (*ET*_0_) estimations by using a machine-learning approach as a means to bridge the accurate lysimeter data and the easier-to-measure environmental data.

### 1.4. Machine-learning approaches

Machine-learning algorithms are designed to handle large amounts of data that are difficult to handle with standard statistical tools. Over the last several years, increasing numbers of studies have utilized machine-learning strategies to improve the prediction of *ET*_0_. This is due to the ability of machine-learning models to identify complex connections and relationships better than empirical equations can, as described by (Ferreira et al., 2022; Manikumari et al., 2020). Those studies have presented *ET*_0_ predictions based on existing calculation methods that incorporate daily means of meteorological parameters, while trying to minimize the number of parameters that are used to calculate the FPME (de Meneses et al., 2020; Ferreira et al., 2019; Kim et al., 2022; Nagappan et al., 2020) and also maintain a high level of accuracy (*R*^2^ > 0.85). However, although those studies involved data collected over a period more than 10 years and from more than 16 meteorological stations, the training of the models was conducted in the first year of measurement and the testing was done in later years, rather than randomly, which raises the question of whether the modern equipment’s resolution and accuracy aided the prediction, by lowering biased data. Furthermore, the use of k-means (clustering) might create a fixed, over-fitted model for the specific area, which is clustered, as compared to a general model approach. Consequently, we question the legitimacy of using this technique to determine the amounts of irrigation to be applied to fields that are relatively distant from the point which measurements are made.

### 1.5. Evaluation of water evaporation using high-resolution measurement data

We hypothesized that greater resolution of collected temporal and spatial data would improve the accuracy of the FPME model. We also hypothesized that using machine-learning algorithms to analyze big data would result in even more accurate predictions of the absolute evaporation, as compared with the actual water-loss rate measured from evaporative baths (using numerous lysimeters).

In this context, we aimed to ascertain whether machine learning that relies on high-resolution meteorological data and actual water-loss data can help us to understand the fluctuations in the microclimate within a greenhouse over the course of the day and throughout the year. We also aimed to investigate the feasibility of enhancing the precision of evaluations of water loss based on data collected from evaporative baths, excluding the impact of biological elements, such as plants. We posit that biological factors could introduce bias into the measurements; therefore, this study seeks to explore the potential for improved accuracy in their absence.

## 2. Materials and methods

### 2.1. Experimental setting

The experiment was conducted in a semi-controlled greenhouse. The greenhouse was exposed to environmental meteorological changes, but cooling pads were used during the summer and a heater was used during cooler periods. The experiment began in December 2020 and ended December 2021. In the greenhouse of the I-CORE Center for Functional Phenotyping (31.904167 N, 34.800654 E), continuous measurements of the evaporation baths and environmental parameters were made using a high-throughput telemetric, gravimetric-based system (Plantarray system; Plant-Ditech, Israel).

### 2.2. Experimental set-up

The experiment included 72 highly sensitive load cells (lysimeters), 62 of which were mounted with an evaporation bath (pan; with a diameter of 22 cm and depth of 5.5 cm). These baths were automatically filled with fresh water every morning at 04:00 (Fig. 1). The other 10 lysimeters were mounted with constant weights to monitor their consistency. The water-evaporation rate and the meteorological conditions were monitored continuously (at 3-min intervals) and simultaneously. Temperature, relative humidity, barometric pressure and VPD were measured using 12 VP-4 meteorological sensors (MS). The water temperatures of 32 evaporating baths were measured using the 5TE sensor. Solar radiation was measured by the greenhouse main meteorological station (MMS) Watchdog 2000 Series weather stations. Wind speed was measured using an ATMOS 22 ultrasonic anemometer. All of these sensors were connected to the Plantarray system, which recorded the data and maintained the auto-filling of the baths.

**Fig. 1.**
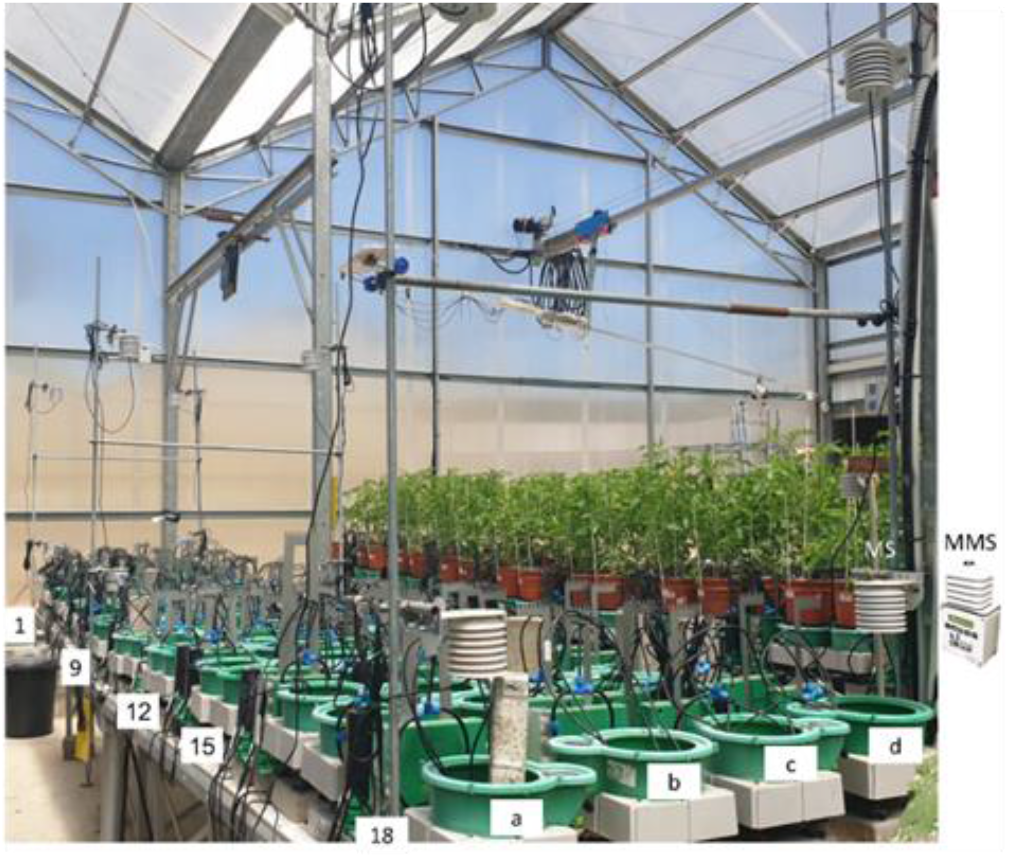
Greenhouse bench equipped with sensors to provide high-resolution data. The experimental bench (7 m long × 1.2 m wide × 0.9 m high) was divided into 18 columns (1–18) and four rows (a–d). A total of 12 meteorological stations (MS) measuring temperature, relative humidity and barometric pressure were positioned at two heights (Z0 = 0.3 m above the baths and Z1 = ∼1.5 m above the baths). On the experimental bench, MS units were placed at the following locations: (1, a), (1, d), (9, a), (9, d), (18, a) and (18, d). The main MS was placed next to the lysimeter at Point 18d.

### 2.3. Equations for environmental modulation

The spatial distribution of temperature, RH, VPD and barometric pressure across the MS units was calculated using the inverse distance weighting (IWD) interpolation. The data were estimated by comparing the actual measurements with interpolated values. The use of three MS units coupled with the IWD formula resulted in very high and significant correlations (*p* < 0.001) for the calculation of temperature and relative humidity at any given point in time or space (see Suppl. Fig. 1).

**Equation 1. Inverse-distance weighting (IWD) interpolation.**

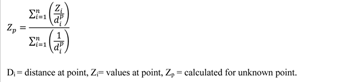

The wind speed in the greenhouse remained constant, averaging 0.324 ms^-1^ throughout the measurement hours. This was achieved by using fans that were turned on every morning before dawn and turned off after sunset.

The FAO-56 Penman-Monteith (FPME) values for temporal (3-min interval) and daily were calculated using Equation 2 based on the measured and calculated meteorological data for each individual location.

**Equation 2. FAO-56 Penman-Monteith equation.**

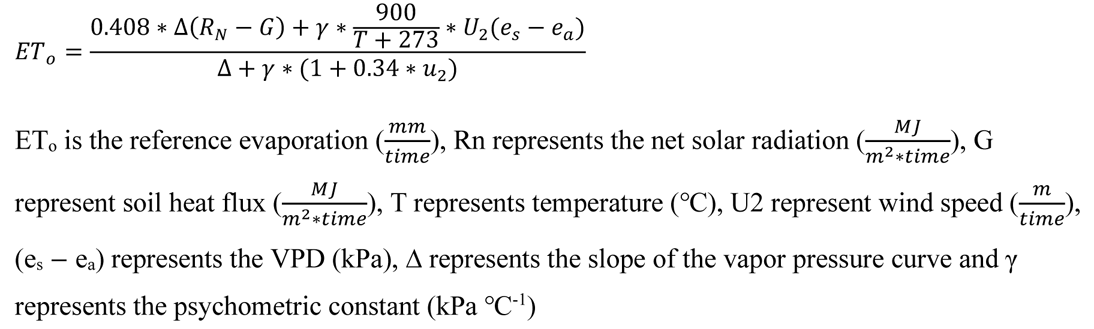

The daily FPME values were calculated based on the interpolated average values of the meteorological factors. As the soil heat flux has a relatively small influence in greenhouses and can be neglected in most situations, it was not included in this analysis.

### 2.4. Machine-learning model and hyperparameters

Several machine-learning algorithms were evaluated, including Decision Tree, Random Forest, XGBoost and Neural Networks. Decision Tree, Random Forest and XGBoost are all example of supervised machine-learning algorithms that are commonly used for classification and regression-prediction models. They are part of a broader family of machine-learning algorithms known as tree-based methods. Those methods are type of nonparametric statistical model that uses a tree-like structure to model relationships between variables in a dataset. In contrast, the Neural Network algorithm is a type of parametric statistical model that uses a set of interconnected nodes (neurons) to model complex relationship between variables.

i. <u>Decision Tree</u>: This algorithm generates a flowchart-like structure (reasonable tree branches), in which each node represents a feature, each branch represents a “decision rule” and each leaf node represents a prediction. The tree is constructed by recursively splitting the given database, with the goal of distinguishing between branches, in order minimize the prediction error.
ii. <u>Random Forest</u>: This algorithm combines multiple decision trees. It creates a collection of decision trees and generates a prediction by averaging the predictions for each individual tree. This approach helps to reduce overfitting (which occurs when a machine-learning model becomes excessively specialized to the training data) and to improve generalization performance.
iii. <u>XGBoost (Extreme Gradient Boosting)</u>: This algorithm sequentially builds an ensemble of prediction models (usually decision trees) and optimizes the objective (gradient-boosting framework) by adding new models that minimize loss. XGBoost adds regularization and advanced optimization, which allow for improved accuracy.
iv. <u>Neural Network:</u> This is a parametric statistical model that uses a set of interconnected nodes (resembling neurons), with complex relationships between the variables. The model consists of an input layer, a hidden layer and an output layer, with each layer containing multiple neurons. Neurons in one layer are connected to the adjacent layer and each connection is evaluated with weighting. The network learns by adjusting the weights based on the training data until it minimizes the prediction error.

The model hyperparameters were set as follows:

i. <u>Decision Tree</u>: The minimum sample split was 2, the minimum leaf sample was 2 and the maximum depth was *none*, meaning that the tree would continue to grow until all of its leaves were pure or until all of its leaves contained less than the minimum sample split sample.
ii. <u>Random Forest</u>: The minimum sample split was 2, the minimum leaf sample was 2 and the maximum depth was *none*. The number of estimators, which is the number of trees in the forest, was set to 100 as a default and bootstrap was set as default (True), meaning samples were used in the building of the trees.
iii. <u>XGBoost</u>: A maximum depth of 100, 55 estimators and a learning rate of 0.3 were selected, which represent the step size used to update model coefficients. Reg_lambda (L2 regularization) was set to 0.3, to prevent overfitting by adding a penalty term to the loss function. Gamma was set to 1e-5, which represents the lowest loss reduction required to further partition the lead node.
iv. <u>Neural Network</u>: The epochs setup was 100, which is the number of iterations over the training dataset. The batch size was set to 1024, which is the number of samples that we needed to work through before updating the model parameters. There were five hidden layers and an Adam optimizer with a learning rate of 1e-3.

To train the models, we filtered out the constant weight places and used the measured and calculated meteorological data and time features (i.e., minute, hour, day week and month) to predict the water weight loss measured by the lysimeters. The training dataset was split, with 70% of the data used for training and the remaining 30% used for testing. For each model, we set up random seed, to expose all of the model to the same training, validation, and test data (several seeds were tested, as we used 42). Based on the absolute error between the predicted and actual results of each machine-learning model, an analysis of variance (ANOVA) was conducted to compare the machine-learning algorithms. That analysis was conducted using JMP 16 Pro statistical software.

## 3. Results

### 3.1. Effects of spatial and temporal variability of meteorological data on evaporation evaluation

The first part of our study focused on investigating the variability of environmental parameters within the greenhouse. To that end, we examined the VPD along with its components (temperature and RH). The data exhibited a wide range of spatial and temporal patterns over the course of the year.

As shown in Figure 2 (a–d), the amount of variability across and between the MS units varied over the course of the year, with greater variability during the warmer seasons. To better understand the cause of this variability, we analyzed the relative contributions of the VPD components, temperature (Fig. 2e–h) and RH (Fig. 2i–l), by calculating the difference between the data collected by the unit at the edge of the bench and the data collected by the main MS. Our result indicats that, across the bench, temperature varied up to 8°C and RH varied up to 16% (THIS WOULD BE CLEARER IF YOU PROVIDED RANGES FOR THE COLLECTED DATA). There was a greater degree of variability in the higher MS, as compared to the lower MS. These results suggest that the variability within the greenhouse was relatively high and that each evaporation bath “sensed” a different microclimate at any given moment. The data from the 12 MS units were used to calculate the meteorological parameters above each of the 62 evaporation baths (pans) using the IWD formula (Equation 1), revealing four-dimensional heterogeneity in the values of the meteorological parameters (Fig. 3).

**Fig. 2.**
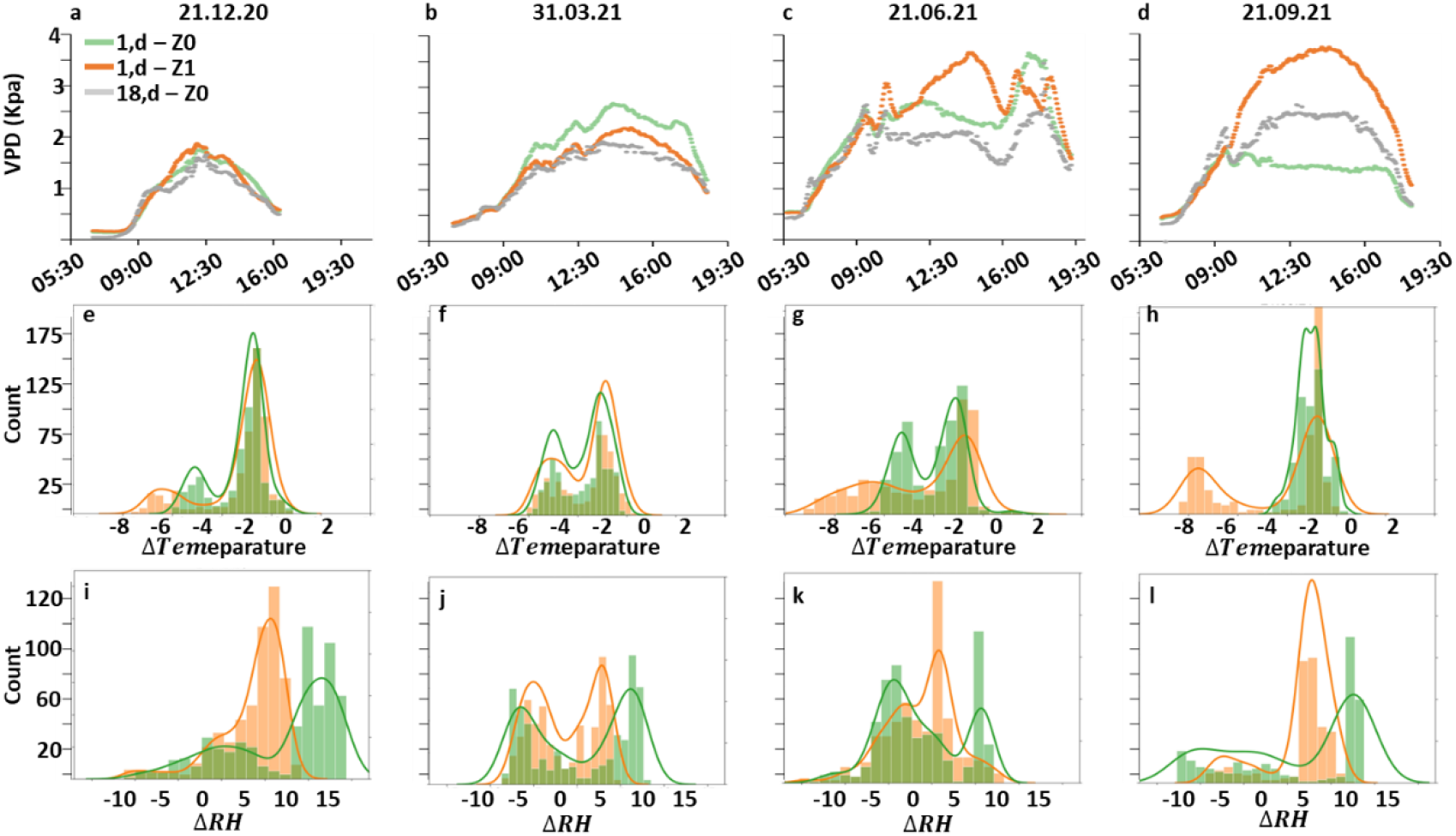
Spatial and temporal variation in VPD above the greenhouse bench. (a–d) Continuous VPD measurements on four key dates. The gray line indicates the data collected by an MS on one side of the bench [Z0 at (18, d)]. Green and orange lines indicate the data collected by the MS units at two heights on the other side of the growth bench [Z0 and Z1 at (1, d)]. (e–g) The change in temperature and (i–l) change in relative humidity across the bench (gradient) over the course of the day for the dates presented above, the “count” on the y-axis represents the number of data points in each interval. The components of the VPD (temperature and RH) were used to determine the disparity among three different spots in the greenhouse: Z0 at (8, d), Z0 at (1, d) and Z1 at (1, d).

**Fig. 3.**
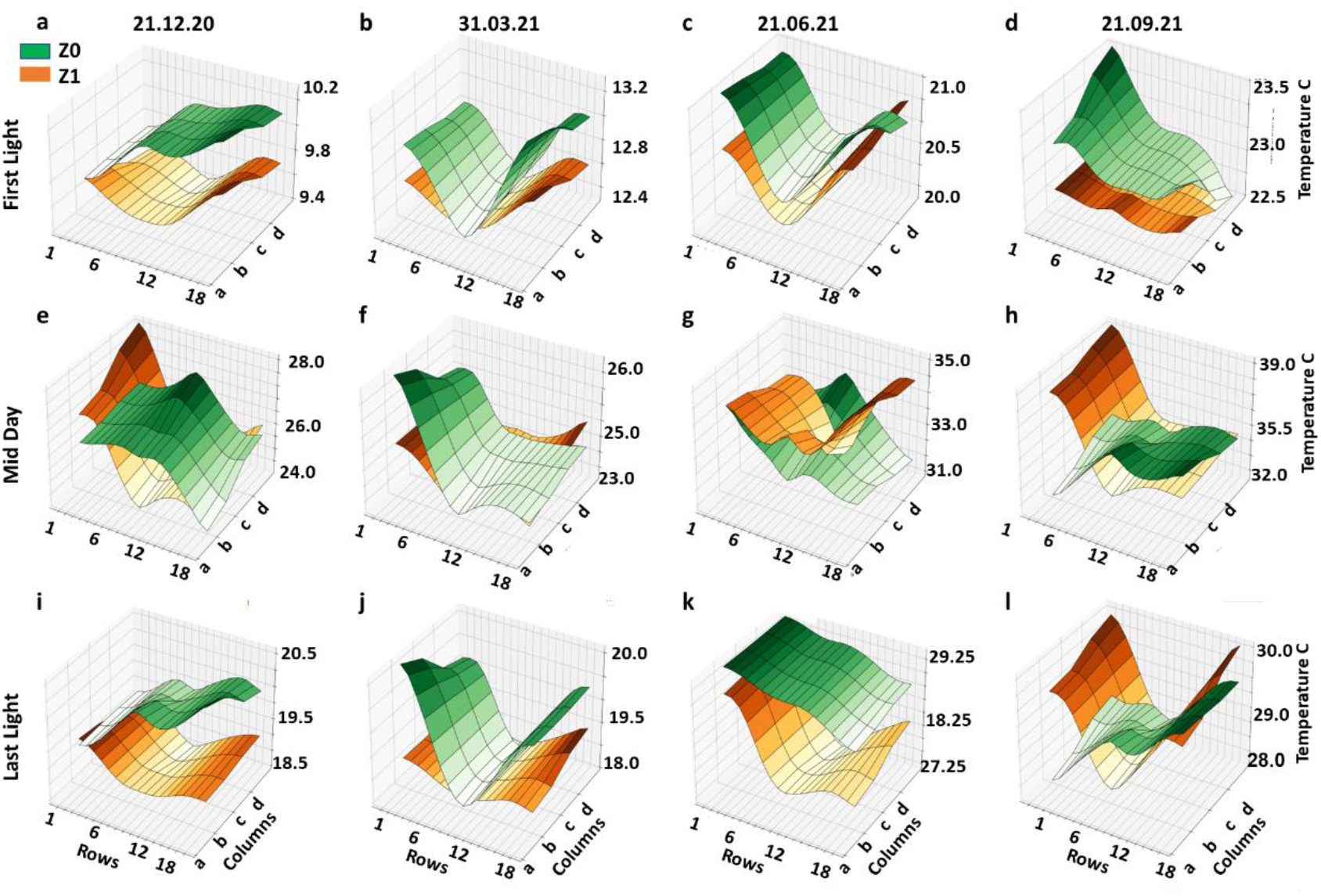
4D projection of temperature over time and space above the 72 load cells on the greenhouse bench. Temperature distribution at Z0 (green) and Z1 (orange) at three times of day on four key dates: (a–d) first solar radiation, (e–h) midday (12:00) and (i–l) last solar radiation. The data for the unmeasured locations were calculated using the IWD formula based on the 12 MS units: six at Z0 and six at Z1 (see Fig. 1, Equation 1). The temperature axis was adjusted according to the minimum and maximum data points for each subplot. The distance between adjacent points was 0.3 m.

The observed variability in temperature over time reveals the unique microclimates above the evaporating baths across the bench and over the course of the day Nevertheless, the actual amount of water that evaporated from the baths, at both Z0 and Z1, did not correlate directly with the VPD (Fig. 4).

**Fig. 4.**
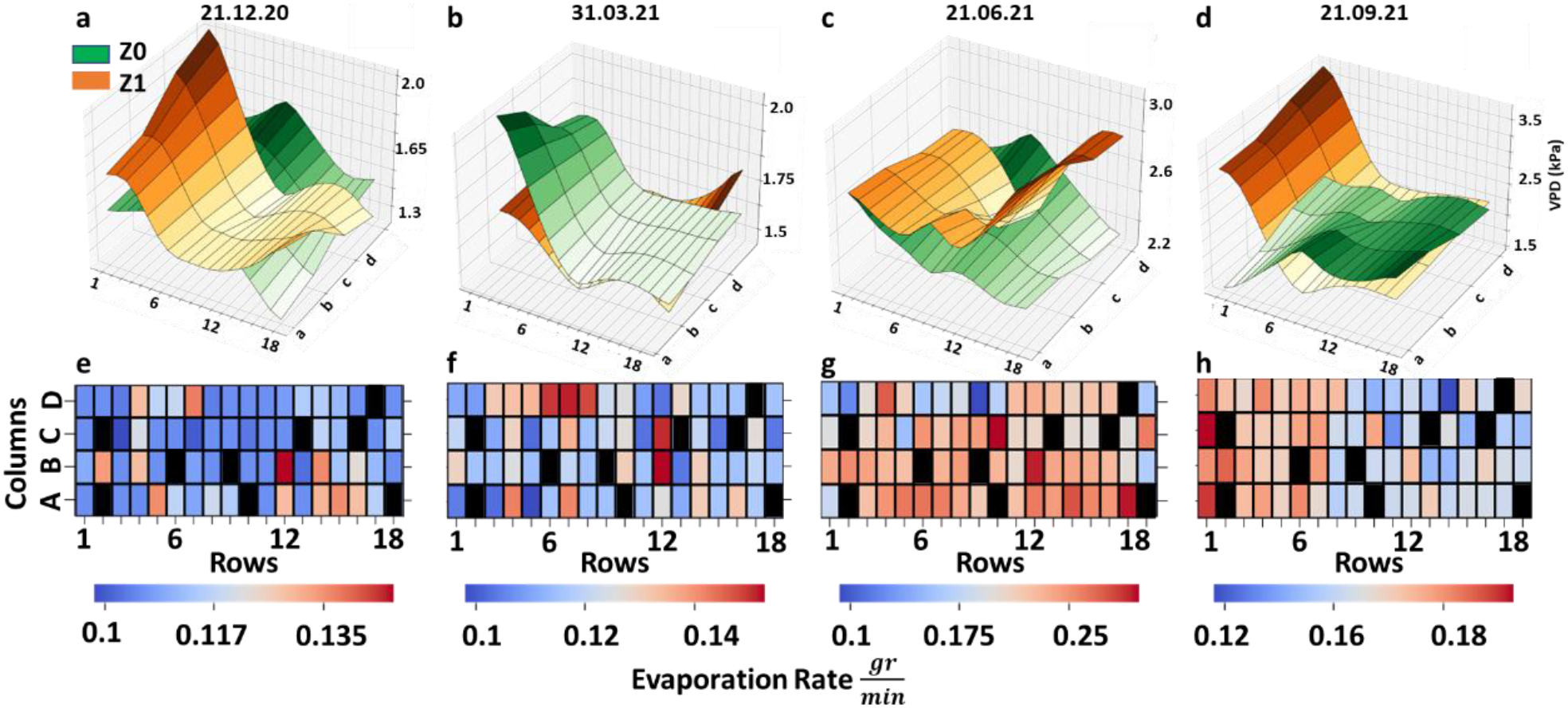
Midday evaporation rate relative to VPD. (a–d) Midday VPD above the baths at Z0 (green) and. Z1 (orange) **on four key dates**. The span of the VPD axis was adjusted according to the minimum and maximum results for each subplot. (e–h) The evaporation rates for 62 water baths as measured using lysimeters. Black zones represent the constant weight scales.

It was surprising to find that neither the VPD measured near the evaporation baths (Z0) nor the VPD measured at a greater height (Z1; Fig. 4a–d) from the evaporation baths had a direct effect on the evaporation rate measured by the lysimeter (Fig. 4 e–h) at midday. This discrepancy between the evaporation rate and the meteorological parameters that lag behind it and were not linearly corelated with one another was revealed to be an example of hysteresis (Fig. 5).

**Fig. 5.**
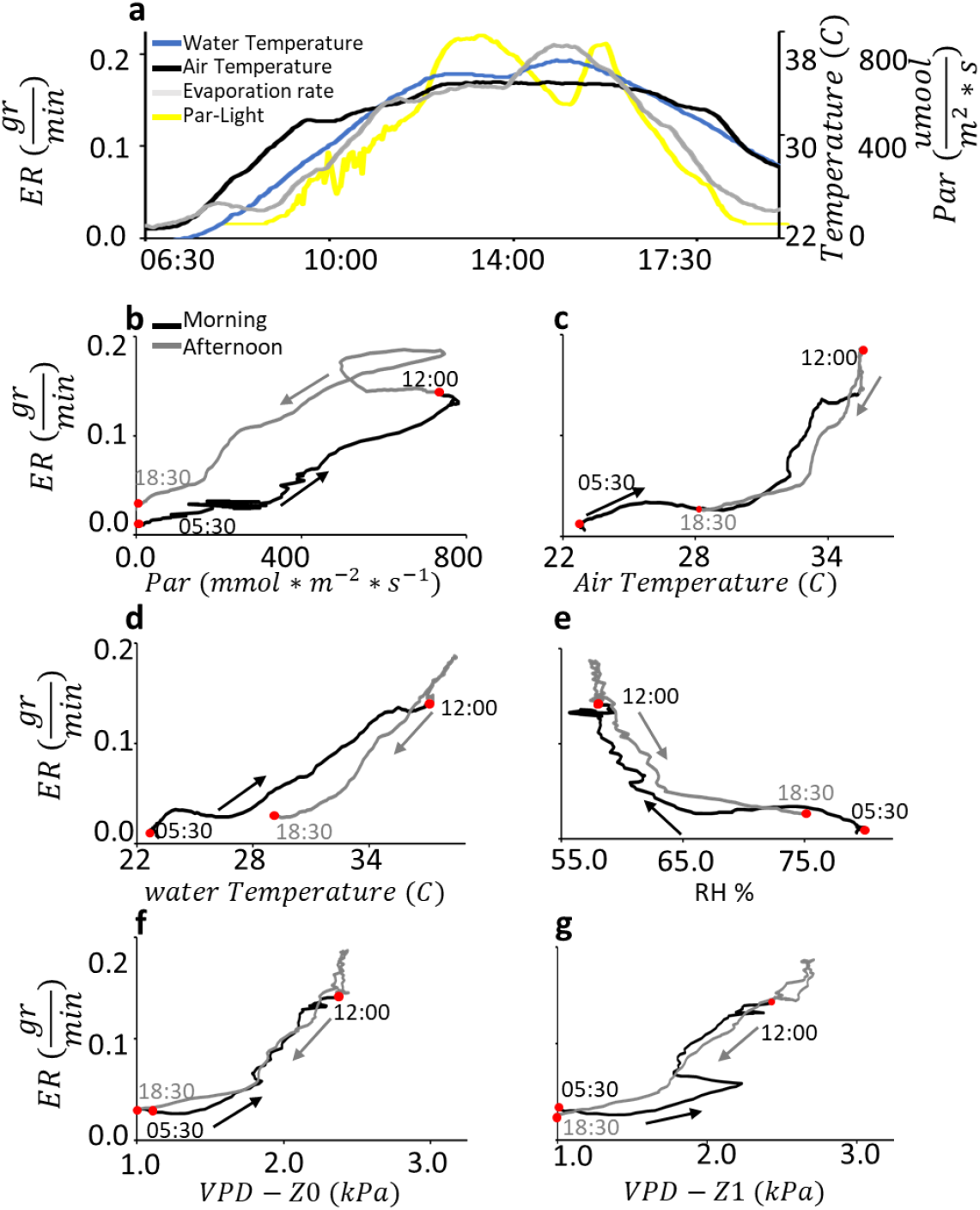
Hysteretic relationships between meteorological parameters and the evaporation rate. (a) Daily patterns of PAR, bath-water temperature, air temperature and the evaporation rate (taken from a representative evaporation bath on 21 Sept. 2021). The temporal correlation between the evaporation rate and (b) PAR, (c) air temperature, (d) water temperature, (e) RH (%), (f) VPD at Z0 and (g) VPD at Z1 revealed nonlinear correlations and hysteretic relationships, as the evaporation rate varied over the course of the day. The changes in the measured data over time are indicated by the black arrows (morning) and the gray arrows (afternoon).

This hysteretic behavior of VPD and the evaporation rate, in which water and air temperature formed a clockwise pattern, while the photosynthetically active radiation (PAR) and RH formed counterclockwise patterns, did not show a specific pattern.

### 3.2. Microclimatic challenges for the FPME model

Those results sharpened our question regarding the accuracy of estimates of evaporation made using models like the FPME. Hysteresis and temporal and spatial variations are taken into account by using calculated average daily values of the meteorological factors values, to test the accuracy of the FPME’s calculations of the evaporation from the baths based on their microclimate conditions. For each bath, we measured the actual absolute daily evaporation (using lysimeters) and compared it with the evaporation values calculated using the IWD formula. We found that the output of the FPME formula differed from the actual measurements in the range of ±22%, depending on the day of the year (Fig. 6).

**Fig. 6.**
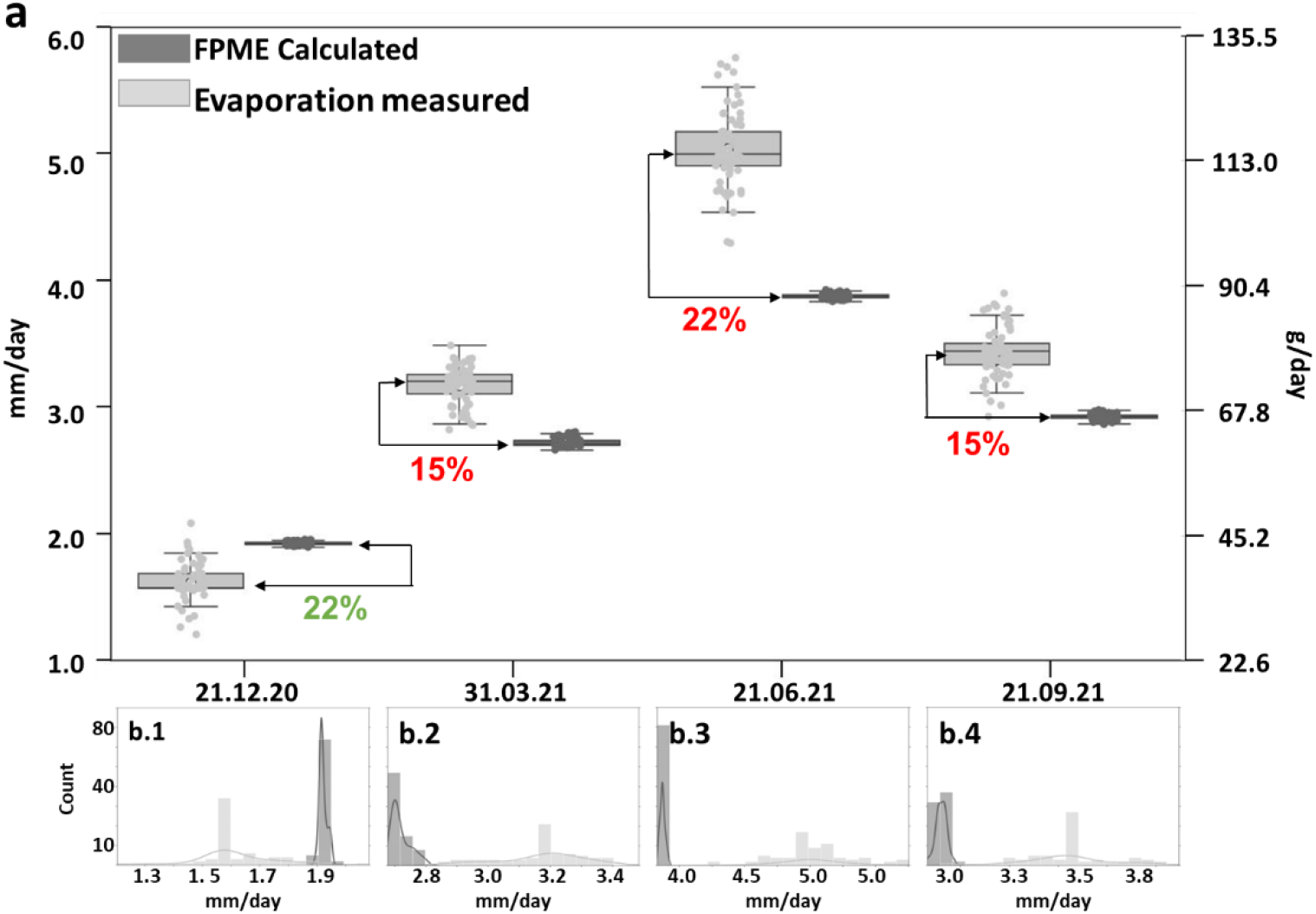
Yearly comparison between the measured daily evaporation and the daily evaporation calculated using the FPME. (a) Each boxplot represents the daily evaporation values for all baths (*n* = 62 for each box), either measured by lysimeters (solar radiation, light gray boxes) or calculated using the FPME (dark gray boxes), on four key dates. The distance between the two horizontal arrows demarcating the percentage values represents the difference between the medians of the calculated and measured values (green text = excess; red text = absence). (b) Daily distribution of the evaporation rates for all baths, as measured by the lysimeters (solar radiation, light gray) and calculated (dark gray) for four key dates.

The low accuracy and denser spread of the FPME-calculated values as compared with the real (unstable) spatial and temporal data led us to wonder whether high-resolution measurements (3-min interval) might improve the accuracy of the FPME. The differences between the momentarily measured and FPME-calculated values were even greater than the differences between the daily measured and FPME-calculated values (Figs. 6 and 7; *R*^2^ = 0.637).

**Fig. 7.**
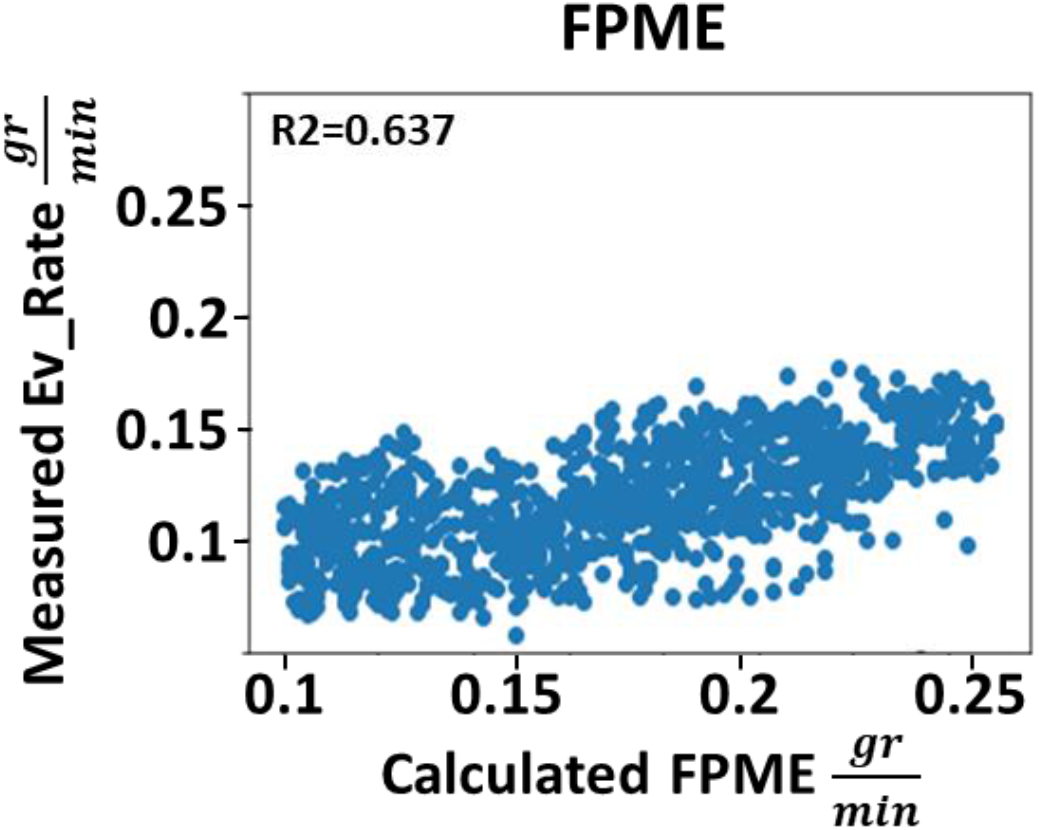
Correlation between the measured evaporation and the evaporation calculated using the FPME, at 3-min intervals. Pearson correlation between the absolute measured evaporation rate (lysimeter-based) and the FPME-based interpolated values. The dataset included 3.01e6 entries (the full dataset).

### Predicting evaporation using machine-learning algorithms

According to our results, the accuracy of the evaporation predictions could be improved if the spatial-temporal variation were taken into account and if higher-resolution measurement data were used. We attempted to use four machine-learning algorithms (Decision Tree, Random Forest, XGBoost and Neural Network). We also used two different types of datasets: direct and indirect. The direct dataset included data for all of the environmental factors, including the bath temperature (assuming that this parameter integrates the ambient parameter with the parameter that has the strongest influence over the evaporation rate). The indirect dataset included data for all of the ambient parameters, except for water temperature.

We were able to predict the evaporation rate within a 3-min interval with a high correlation to the measured data using all four models (*R*^2^ values ranged between 0.918 and 0.976). Interestingly, adding the direct measurements did not significantly improve the models’ accuracy, but the XGBoost model performed significantly better than the other models (Fig. 8e; Indirect + Direct). All model types and the use of both direct and indirect + direct measurements provided better predictions than the FPME (Fig. 7).

**Fig. 8.**
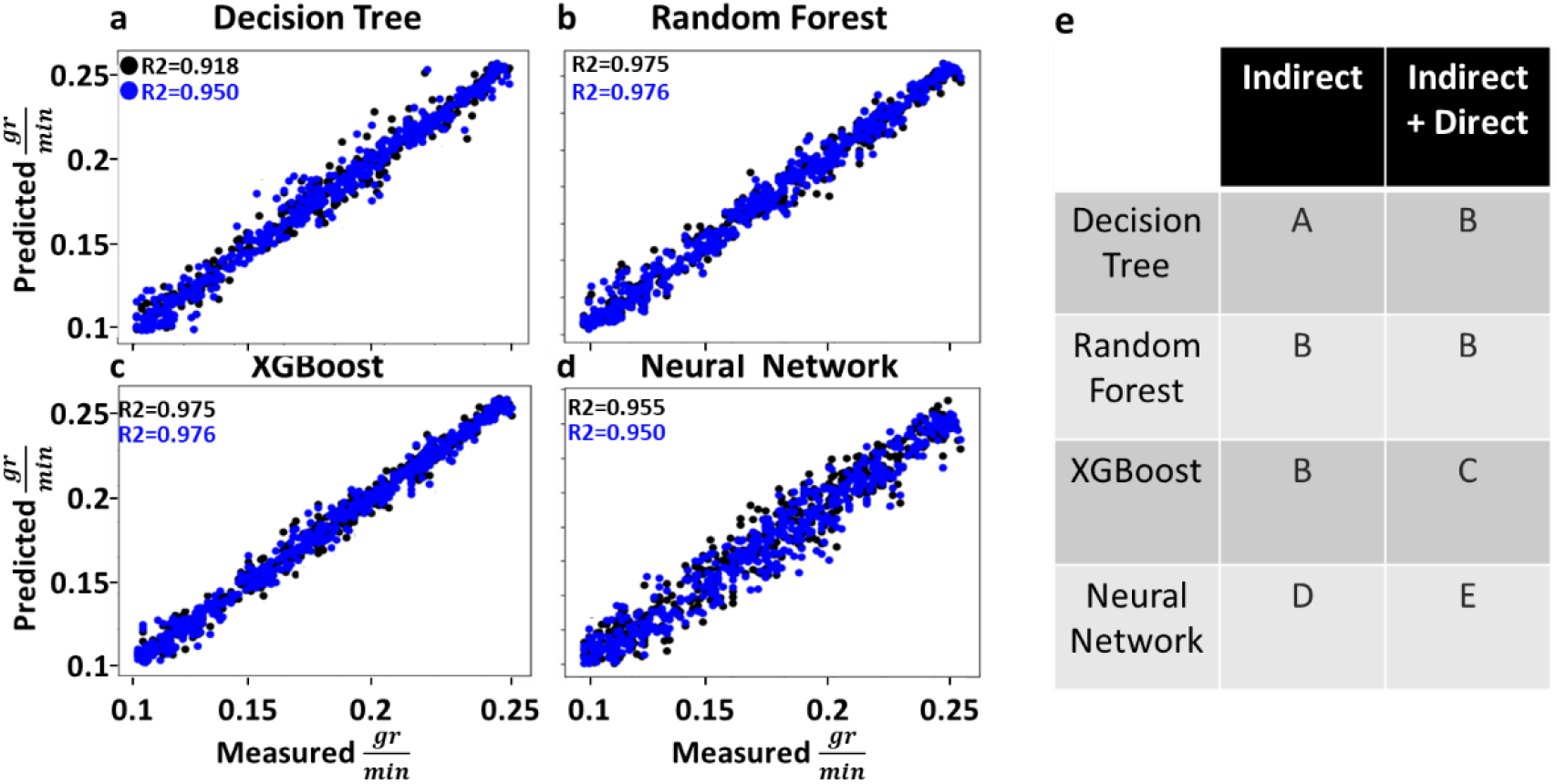
Evaluation of the performance of the different algorithms based on direct and indirect + direct datasets. A dataset including 1.2e6 data points describing meteorological parameters and bath-water temperature was used to evaluate the evaporation rate. A model was separately tested for both a dataset that did not include bath-water temperature (indirect; black dots) and a dataset that did include bath-water temperature (indirect + direct; blue dots). The four algorithms were all applied to the same dataset (70% train, 30% test): (a) Decision Tree, (b) Random Forest, (c) XGBoost and (d) Neural Network. (e) Different letters indicate a significant difference (*p* < 0.001), according to ANOVA of the absolute error of each individual test (indirect and indirect + direct models).

To identify the key input features that have the greatest impact on the models’ predictions, feature-importance techniques were used to assign scores to the input features (Li et al., 2017). These scores offer insights into the relative significance of each input feature and can help us to understand the hierarchical relationships among them (Fig. 9).

**Fig. 9.**
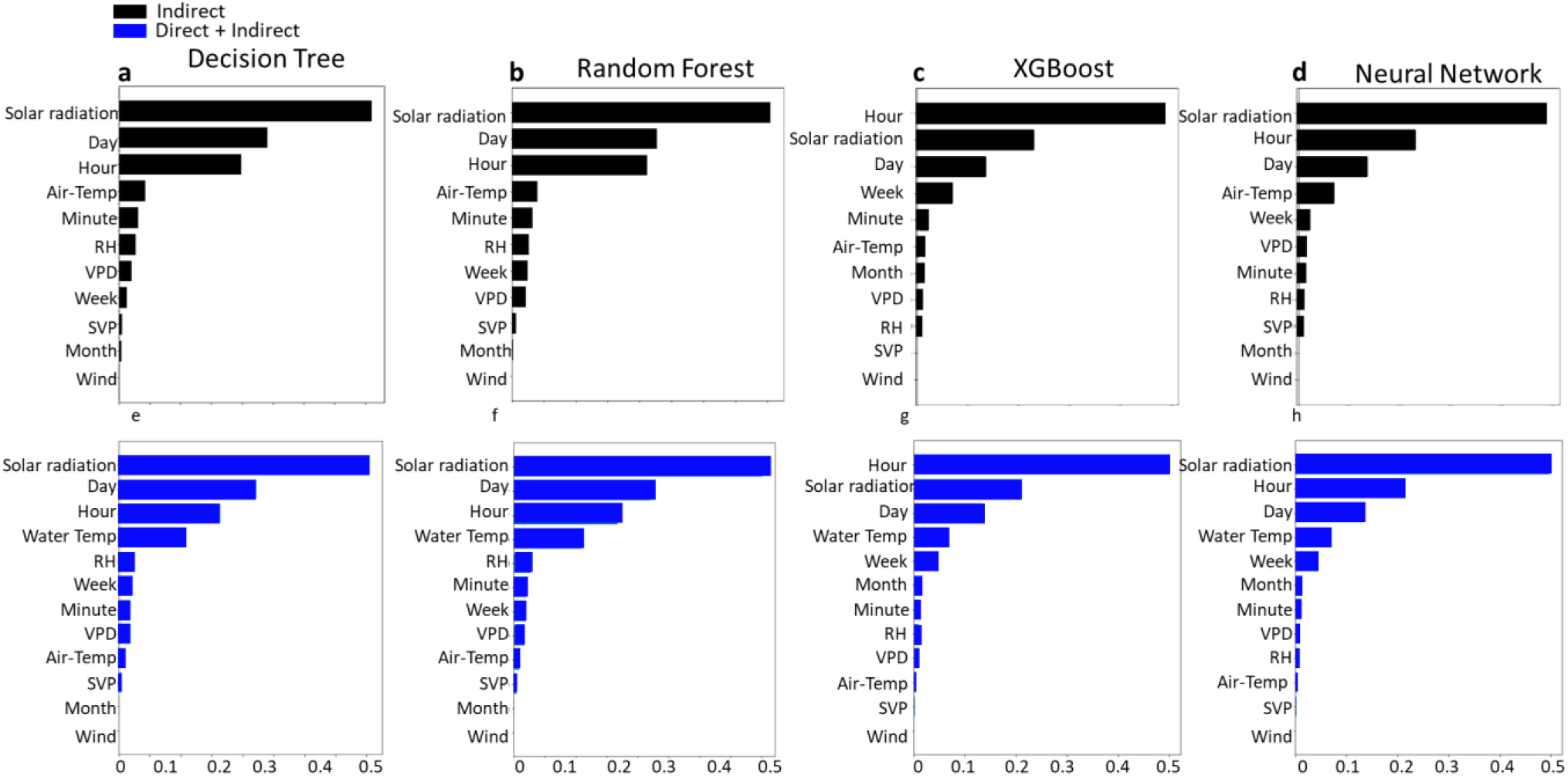
Indirect and indirect + direct model score-importance hierarchy. Each plot represents the feature hierarchy for each of the algorithms, indirect (black) and indirect + direct (blue).

Solar radiation was the top feature in all of the (direct and indirect) models, except for XGBoost (Fig. 9c,g), in which hour was the feature with the highest impact score, followed by solar radiation. XGBoost’s unique hierarchy also resulted in the highest *R*^2^value (0.976). The third most influential factor in all of the direct and indirect models was also time-based (i.e., day or hour). The fourth most important feature was the first to differ between the direct and indirect models. In all of the direct models, that fourth feature was bath temperature (Fig. 9e,f,g,h). However, air temperature was the fourth most important feature in all but the XGBoost model (Figs. 9a,b,d), which implies that the heating of water by the air makes a greater contribution to evaporation than air temperature (or VPD, in this instance).

To better understand the role of water temperature, we re-trained our model based on water temperature and the temporal feature alone (Fig. 10).

**Fig. 10.**
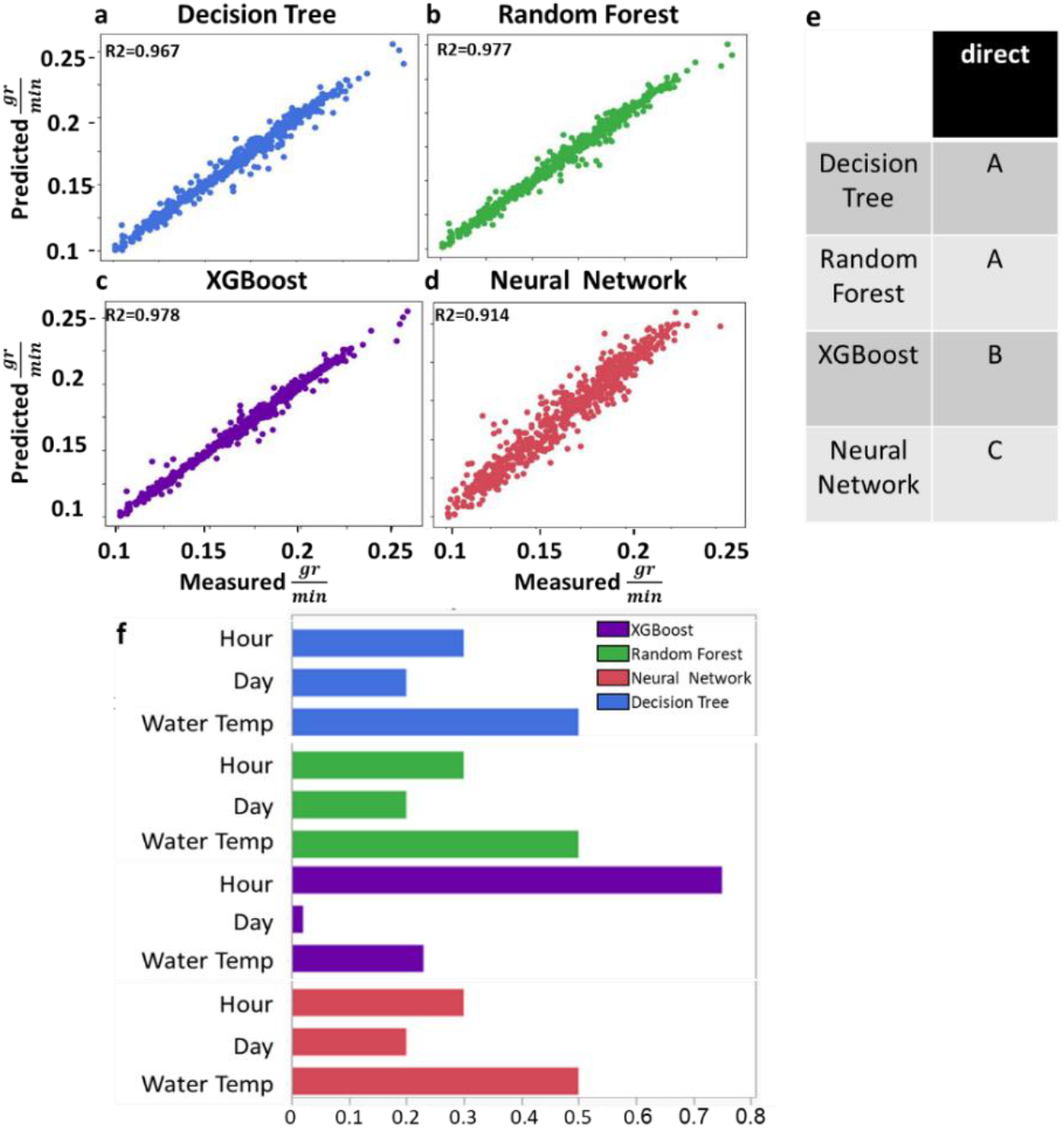
Comparison of the machine-learning models based on water temperature and temporal data. A dataset including 1.2e6 data points for bath temperature and temporal parameters was used to compare the measured values with those predicted by the different algorithms. All of the algorithms were applied to the same dataset (70% train, 30% test). (a) Decision Tree, (b) Random Forest, (c) XGBoost and (d) Neural Network. (e) Connecting-letter report based on ANOVA of the absolute error of each individual test (indirect and indirect + direct models). Different letters indicate a significant difference (*p* < 0.001). (f) Feature hierarchies of the different algorithms: Decision Tree (blue), Random Forest (green), XGBoost (purple) and Neural Network (red).

The direct models based on bath temperature, hour and day performed better across all of the different types of algorithms, as compared with the all-feature model (*R*^2^values ranged between 0.914 and 0.978). XGBoost was the most accurate, followed by Random Forest, Decision Tree, and Neural Network. A connecting-letter report (Fig. 10e) revealed a result that was similar to that obtained using the larger dataset (Fig. 8e). With regard to Decision Tree, Random Forest and Neural Network, the water temperature, hour, and day features appeared in the order of the direct measurement features. The XGBoost model emphasized hour as the most important factor, followed by day and water temperature. According to the connecting-letter report, XGBoost models accuracy was significantly higher compared to the other model However, in those models, water temperature was only the fourth most important feature (see Suppl. Fig. 2).

### 3.3. Temporal and spatial limitations of the machine-learning algorithms

The accuracy of a time-series model is influenced by the time-interval resolution and the total amount of data; as the quantity of data increases, the model should become more accurate (Es-Sabery et al., 2021). To determine the accuracy of our models, we tested the impact of both time interval (increased by 3-min collection resolution) and the total amount of data (Fig. 11).

**Fig. 11.**
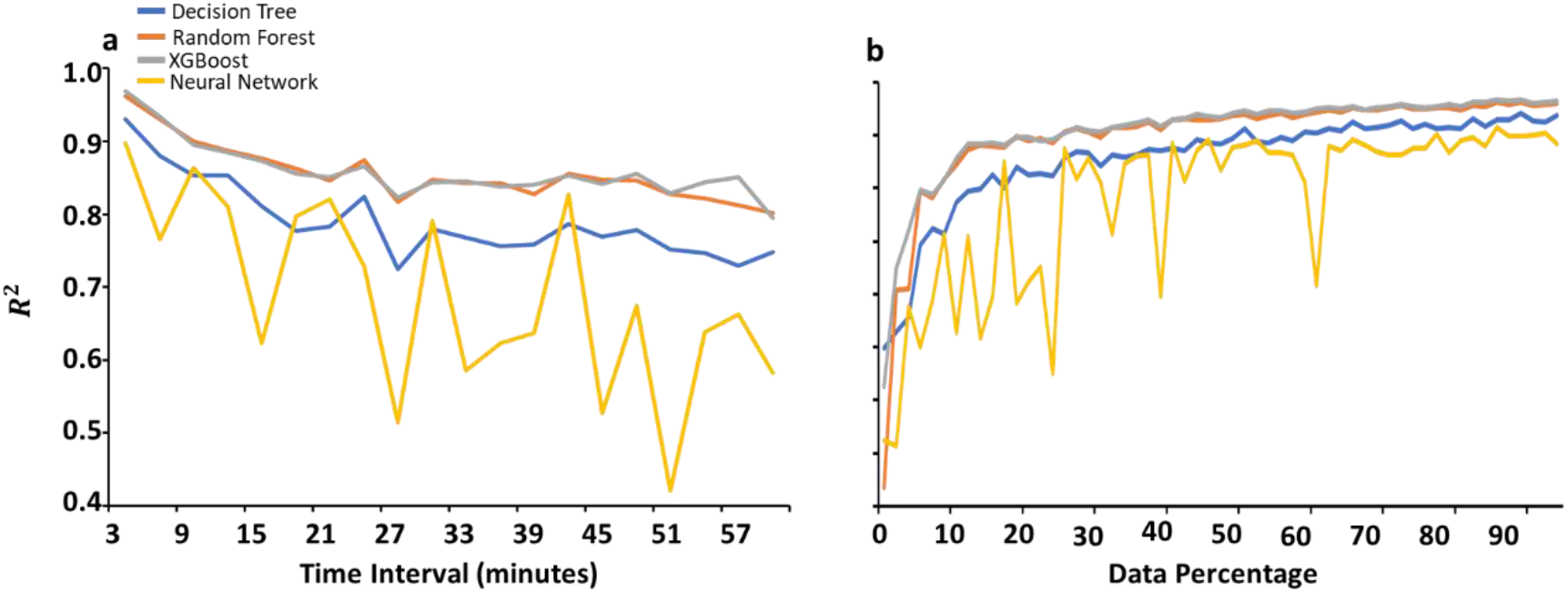
The influence of data resolution and data quantity on the accuracy of the prediction models. *R*2 values for. Decision Tree (blue), Random Forest (orange), XGBoost (gray) and Neural Network (yellow) (a) when data were collected at different time intervals and (b) when different proportions of the set of directly measured data were used, with each data sample randomized.

As the time interval between measurements increased, the accuracy score for each model decreased. In all models (Fig. 11a), both Random Forest and XGBoost generated similar *R*^2^scores and stabilized at ∼30-min sampling rate with a *R*^2^of 0.84. In contrast, the *R*^2^ value for the Decision Trees and the same sampling rate was 0.72. Interestingly, as the time intervals were increased, the Neural Network *R*^2^ fluctuated without any noticeable stabilization. As the proportion of data used increased, the accuracy scores for all of the models increased in a logarithmic pattern (Fig. 11b). Yet, while Random Forest, XGBoost and Decision Tree reached near-stabilization after 20–30% of the total data had been incorporated into the model (*R*^2^ values of 0.907, 0.908 and 0.867, respectively), the Neural Network values fluctuated until ∼65% of the data had been included (*R*^2^ = ∼0.87).

That finding suggests that 20–30% of the original temporal dataset is sufficient for obtaining highly accurate predictions (*R*^2^ > 0.85) from all algorithms, except for Neural Network. To test that assumption, we used 10% of the original dataset (one full month of data) and examined the above models’ *R*^2^ scores (Fig. 12).

**Fig. 12.**
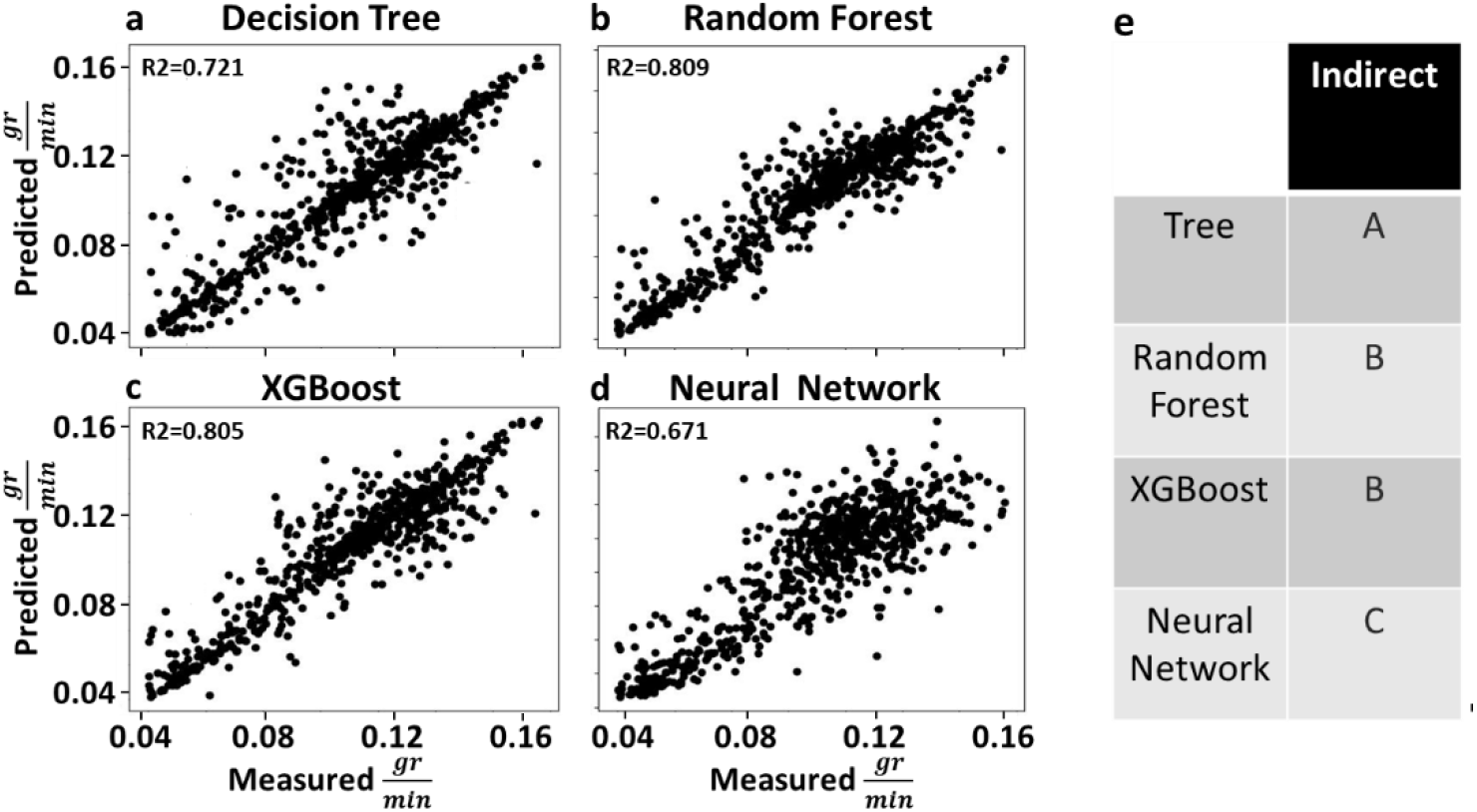
Comparison of algorithms using 10% of the temporal dataset. This dataset included 3.01e5 data points for meteorological parameters (10% of the original dataset). Only the indirect-measurement data were compared with the actual measured data. Each algorithm was applied to the same dataset (70% train, 30% test). (a) Decision Tree, (b) Random Forest, (c) XGBoost and (d) Neural Network. (e) Connecting-letter report based on ANOVA of the absolute error of each individual test (indirect and indirect + direct models). Different letters indicate a significant difference (*p* < 0.001).

When 10% of the data were used, Random Forest and XGBoost lost 0.02 of the variation in the prediction accuracy, Decision Tree lost 0.046 and Neural Network lost 0.226 (*R*^2^ = 0.913, 0.935, 0.861 and 0.696, respectively), as compared with the full-dataset model (Fig. 8). Thus far, our results indicate that approximately 21.5e3 environmental data points collected at 3-min intervals can be used to accurately predict the rate at which water will evaporate from a bath (Fig. 11b). To confirm these results, we performed an similar independent experiment in a different area (the opposite side of the original bench) in the greenhouse, with fewer data points and greater spatial differences. In that experiment, we used only three evaporation baths, which were positioned ∼2.5 m apart from each other, and collected data at 3-min intervals for 1 month (Fig. 13).

**Fig. 13.**
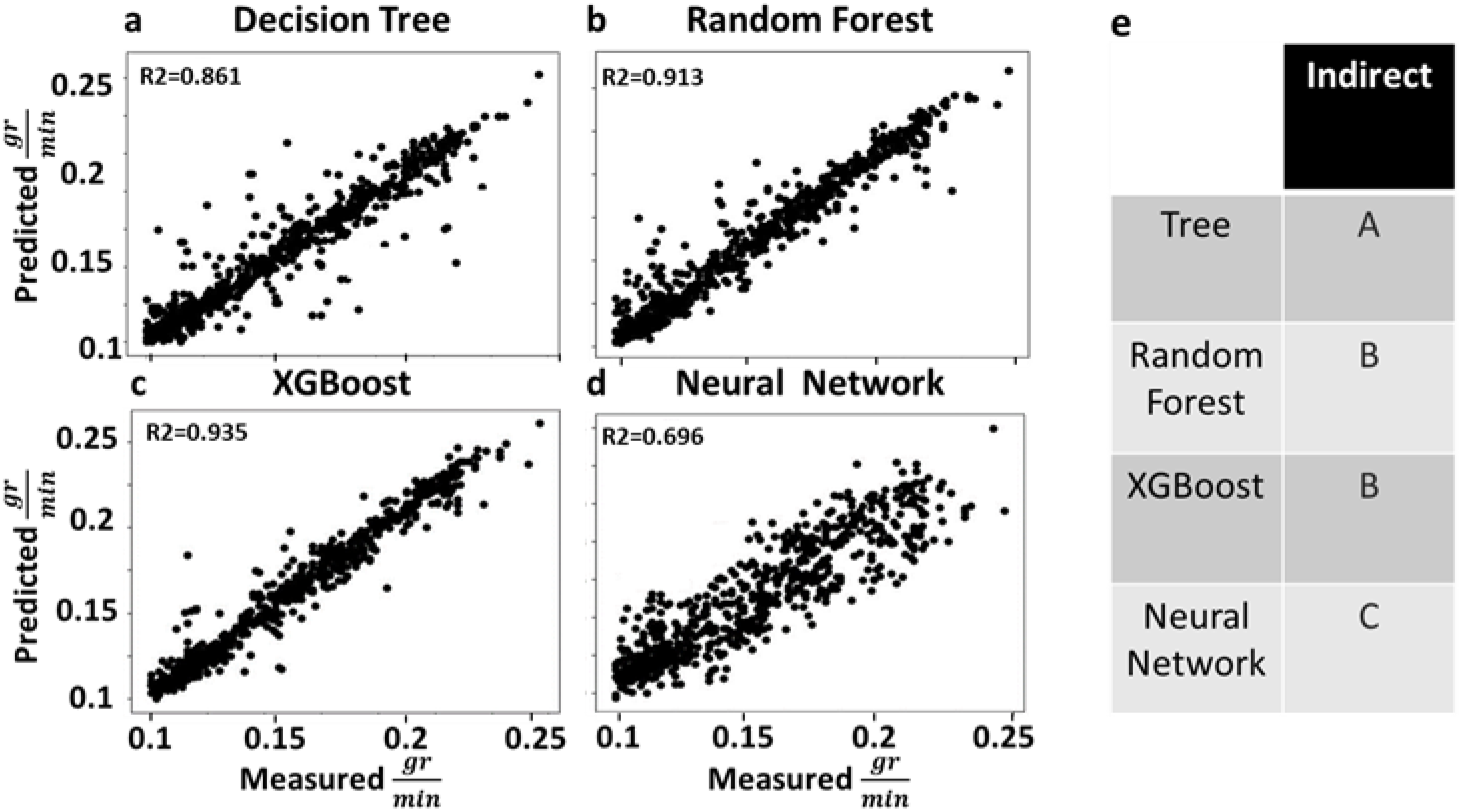
The performance of the different algorithms when a smaller space and a smaller dataset were used. The dataset included 21.5e3 data points for meteorological parameters (∼0.715% of the first experiment), including indirect measurement data used to evaluate the evaporation rate. Each algorithm was applied to the same dataset (70% train, 30% test). (a) Decision Tree, (b) Random Forest, (c) XGBoost and (d) Neural Network. For each of the algorithms, the correlation between the measured and predicted evaporation rate was <0.001. (e) Connecting-letter report based on ANOVA of the absolute error of each individual test (indirect and indirect + direct model). Different letters indicate a significant difference (*p* < 0.001).

Taking substantially fewer measurements and considering larger spatial differences across the experiment bench, solar radiation lowered the accuracy of the predictions produced by the algorithms (*R*^2^= 0.809, 0.805 0.721 and 0.671, respectively), as compared to the full and the 10% datasets (Fig. 8 and Fig. 12 respectively). Nevertheless, this approach still provided satisfactory results, with the accuracy of all of the algorithms’ considerations of the spatial feature receiving high *R*^2^values, as compared to the FPME score (Fig. 7). To gain further confirmation regarding the spatial magnitude of our data, we re-trained our dataset by considering both direct measurements from the experiment locations (Fig. 8, Fig. 13) and samples from three evaporating baths on each bench (Fig. 14). The evaporation baths were distributed across the area, with Dataset 1 (Fig. 8) including the locations (2, a), (9, d) and (18, c) and Dataset 2 (Fig. 13) including the locations (1, d), (9, a) and (18, d).

**Fig. 14.**
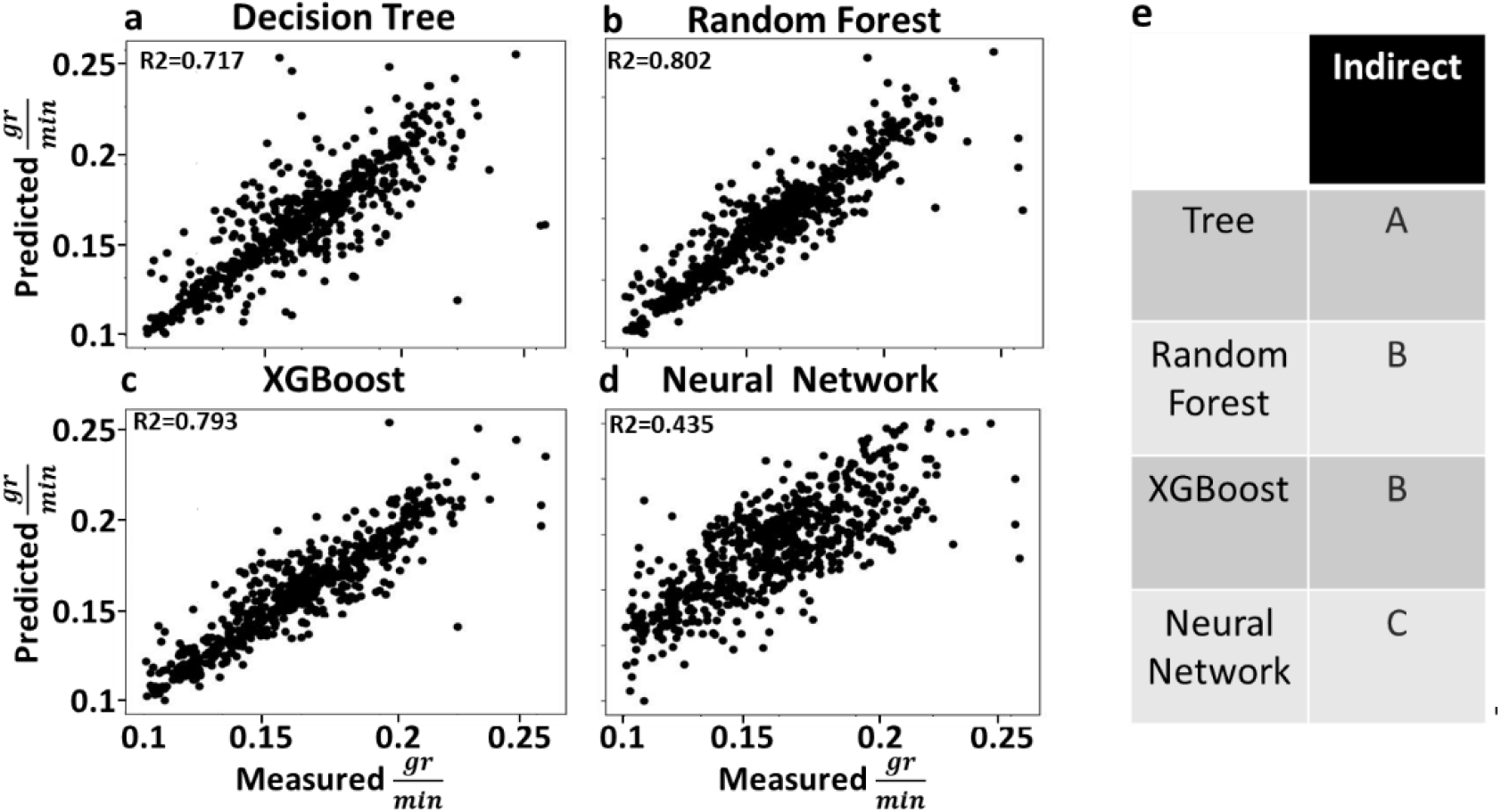
Comparison between machine-learning models applied at both locations. The dataset included 36.806e3 data points for the meteorological parameters (∼1.22% of the dataset used for first experiment), including indirect measurements to evaluate the evaporation rate. Each model was applied to the same dataset (70% train, 30% test). (a) Decision Tree, (b) Random Forest, (c) XGBoost and (d) Neural Network. (For each of the algorithms, the correlation between the measured and predicted evaporation rate was <0.001. (e) Connecting-letter report based on ANOVA of the absolute error of each individual test (indirect and indirect + direct models). Different letters indicate a significant difference (*p* < 0.001).

With the expansion of the dataset and spatial range across two edges of the greenhouse, the accuracy of the machine-learning algorithms decreased. In comparison to the full dataset, the 10% dataset and the independent experiment, the accuracy of the Neural Network results declined markedly (*R*^2^ = 0.717, 0.802, 0793 and 0.435). Nevertheless, all of the algorithms, except for the Neural Network algorithm, yielded better results than the FPME (Fig. 8).

## 4. Discussion

In this study, we evaluated whether it is possible to improve the accuracy of the FPME prediction model by taking into account temporal and spatial heterogeneity in the greenhouse. We found that, contrary to our hypothesis, feeding FPME with data regarding ambient conditions across greater temporal and spatial ranges resulted in less accurate predictions of evaporation. In contrast, our second hypothesis was proven to be correct, as machine-learning algorithms using the same big data were able to predict evaporation based on ambient conditions with significantly greater precision than the FPME. We attribute these differences to the fact that the FPME averages values over longer periods (such as whole days or weeks), thereby neglecting cumulative time factors. As a result, it diminishes the impact of daily hysteresis events, potentially leading to inaccuracies. On the other hand, machine-learning algorithms (i.e., Decision Tree, Random Forest, XGBoost and Neural Network) can handle nonlinear relationships in the data, allowing for the consideration of hysteresis factors and achieving much greater accuracy.

### 4.1. Microclimatic challenges to the FPME model

The spatial and temporal heterogeneity of meteorological parameters across the experiment bench (Fig. 2) supported our basic research rationale regarding the heterogeneity of the greenhouse agronomic environment, with each bath exposed to different, dynamic microclimatic conditions. This heterogeneity is a result of the physical structure of the greenhouse (e.g., top beams shade, side walls warm up, etc.), as presented previously (Nebbali et al., 2012) and is expected to be an issue in any growth facility. Such microclimatic heterogeneity necessitates the use of many sensors, in order to get reliable spatial resolution. Our results revealed that the use of the IWD formula (Equation 1) and a relatively small number of sensors is sufficient to provide an accurate 4D presentation of the ambient conditions (Fig. 3), with three sensors sufficient to calculate an accurate assessment point between them (Suppl. Fig. 1; *p* < 0.001). This allows us to use many more spatial data points.

Nevertheless, despite the high resolution of the spatial and temporal data regarding the ambient conditions in the greenhouse, we were not able to improve the accuracy of the FPME predictions of the daily (error range of ±22% ; Fig. 6) or the momentary evaporation (*R*^2^= 0.637; Fig. 7). Furthermore, at the daily scale, FPME predicted evaporation values that were much more homogenous and more narrowly distributed than the measured evaporation values (Fig. 6b). This suggests that there is some smoothing of the data in the FPME model. We suggest that this FPME narrow-distribution disparity is due to the averaging meteorological values, which reduces the magnitude of the effects of major events during the measurement period (Batalha et al., 2018), resulting in a narrower distribution. We suggest that the reduction in the accuracy of the FPME’s momentary predictions, despite the increased number of data points is a result of the hysteretic impact of the evaporation rate not responding immediately to ambient meteorological conditions (Fig. 5). As water heat storage increases or decreases over the course of the day, the evaporation rate will react accordingly due to the changes in phase from water to gas molecules (McCuen & Asmussen, 2009).

Indeed, we demonstrated the hysteresis of different meteorological parameters over the course of the day, each of which has different-sized effects on the evaporation rate (Fig. 5), as (Cui et al., 2016) has previously demonstrated for large water compounds. As other studies have demonstrated previously, the linearization of the FPME results in deviation, which introduces errors into the estimation and, therefore, a non-linear solution should be considered (Leca et al., 2011; Moran et al., 1996; Paw U, 1992). Moreover, as described by (Widmoser, 2009), the discrepancy in estimations varies between daily and hourly rates, with larger differences observed for the latter. This finding aligns with the results that we present in Figures 6 and 7.

Therefore, our hypothesis that averaging the daily parameters would affect the accuracy of the daily FPME prediction was found to be related to the disregarding of the magnitude of the hysteresis. We know that the momentary FPME predictions do not take into account temporal changes (hysteresis), which may explain their inaccuracy. Moreover, this averaging also explains why the daily FPME is relatively more accurate than the momentary one, as averaging daily hysteresis reduces the momentary noise.

The independent complexity of the multi-level, multi-factor variable affecting the evaporation rate makes it challenging to evaluate the evaporation rate. To address this complexity, we used machine-learning algorithms that are designed to take into account complex and unordered changes, in an effort to predict evaporation rates more accurately.

### 4.2. Using machine-learning algorithms to predict evaporation under multi-level, multi-factor, hysteretic ambient conditions

In this work, machine-learning algorithms showed outstanding results when trained with all indirect and direct ambient conditions. Interestingly, the addition of the direct factor of water temperature (Fig. 8a–d; blue dots) improved the performance of all of the tree-based machine-learning algorithms, but had a negative effect on the performance of the Neural Network algorithm.

To better understand the output of the different algorithms, we decided to take a closer look at their feature components and how they are linked to our hypothesis. Many people consider machine-learning algorithms to be a black boxes (Azodi et al., 2020), due to the complexity of the consideration of multiple parameters and statistical calculations of the models, which make it hard to understand the dependency relations between multiple variables. However, utilizing the algorithms’ hierarchical feature (Fig. 9) allows for a better understanding of the relative strength of the effects of the different features on the algorithms’ predictions.

In all of the algorithms except for XGBoost, solar radiation was the most important feature affecting evaporation. This could be due to the redundancy in our dataset. Our solar-radiation measurement relied on a single solar-radiation sensor and assumed the homogeneous distribution of solar radiation throughout the greenhouse. Consequently, this led to a high level of repetition of identical solar radiation measurements for each timestamp entry, potentially biasing the model toward this value and overemphasizing its significance. Furthermore, all of our models suggested that wind speed was the least important feature. That might be due to the fact that we applied a constant wind speed that can be considered relatively low and can be neglected (Varga-Haszonits et al., 2022). The combination of low wind speed and the size of the greenhouse could lead to a decoupled microclimate. In such a scenario, radiative parameters have a greater impact on evaporation, as previously demonstrated in pepper and banana greenhouses screenhouses (Yohanani et al., 2022).

The hierarchy feature also supports the fact that the magnitude of hysteresis during the day has to be taken into account as a major factor affecting the evaporation rate. The most important indication of that is the fact that, in all of the algorithms, the short-term temporal (day and hour) parameters were among the top three influential features (Fig. 9). While the hour factor was the most important feature in the most accurate model, the XGBoost model (Fig. 9c,e; indirect and indirect + direct, respectively), it was positioned even higher in the hierarchy than the most direct factor, bath water temperature, which was the fourth most important feature (Fig. 9e– h). This confirms the importance of the hysteretic heat storage of the water, namely, its delayed impact on evaporation (i.e., it took some time to translate the air temperature into water temperature that led to the evaporation of water from the bath).

Indeed, using only water temperature and temporal features (without any other environmental parameters) improved the performance of XGBoost, to provide the most accurate result (*R*^2^= 0.978). Yet again, this model suggests that hour is the most important feature (Fig. 10). That finding further underscores the importance of hysteresis in evaporation prediction. Nevertheless, we find it hard to explain why water temperature was only the fourth most important feature, although physically it was thought to be the most direct factor in the model.

### 4.3. Temporal limitations of the machine-learning algorithms for predicting evaporation

Additional support for the importance of high-resolution temporal data was revealed when we examined the contribution of the time measurement interval to the accuracy of the predictions produced by each algorithm (Fig. 11a). These results indicate that a faster monitoring rate (momentary sampling rate) will have an important role in the prediction down to about a half-hour window. Therefore, the sample should be taken at the smallest timestamp, ideally at least 2 times faster than the phenomenon being measured, following the Nyquist frequency principle (Detwiler et al., 2023).

The amount of data also plays a major role (Oates & Jensen, 1997) in the accuracy of the machine-learning methods (Fig. 11b). Neural Network showed a high level of fluctuation until ∼80% (total of 2.41e6 data points) of the data were used, which might explain the decrease in the accuracy of the direct model (Fig. 12d) when the amount of data decreased. However, all of the other models exhibited quite high accuracy while using ∼10% (total of 3.01e5 data points) fully randomized sample datapoints of the full indirect dataset (Fig. 12). These results are consistent with those of other studies that have demonstrated that tree-based machine-learning algorithms perform better than neural networks until a certain amount of data has been entered, at which point neural networks are more advantageous (Kraus et al., 2020; Tang et al., 2018).

Our results suggest that spatial measurement has a major effect on the accuracy of prediction models. Therefore, we conclude that using a high sample rate to account for the heterogeneity of the microclimate within the measurement area is a useful strategy even for relatively shorter periods. Indeed, the results of a new, independent experiment conducted in the other part of the greenhouse, using only three bathss and extending over only 1 month (Fig. 13), showed some similarities to the previous results, with Random Forest having the greatest accuracy followed by XGBoost, although there was no evidence to suggest any significant difference between those algorithms. In line with our previous result, both of those models performed significantly better than the other models and the accuracy of the Neural Network model was decreased by the smaller number of data points. Furthermore, all of the models out-performed the FPME. Thus, this independent modified dataset experiment helps us to conclude that a smaller dataset (21.5e3; 0.715% of the full dataset) based on a shorter measurement period, considering temporal and spatial differences, can yield accurate predictions. Furthermore, the fact that data from only three evaporation baths monitored over 1 month and spread across the greenhouse benches yielded highly accurate predictions proves the importance of spatial measurement.

### 4.4. Spatial limitations of the machine-learning algorithms in predicting evaporation

We found it quite bizarre that the use of the trained models of the full dataset (Fig. 8; black line) did not provide valid predictions (fit valid accuracy of our direct measurement; see Suppl. Fig. 3) of the new set of data collected on the other side of the greenhouse (Fig. 13). We hypothesize that variations in the spatial positions of the water baths led to distinct thermal energy retention responses, influenced by the heterogeneous spatial and temporal microclimatic conditions experienced throughout the day.

To test that hypothesis, we re-trained our model with both benches, on the east and west ends of the greenhouse (Fig. 14). The model result confirmed our hypothesis regarding the magnitude of the effect of the spatial distribution of measurements, in comparison to the amount of data gathered. Random Forest remained the most accurate method, followed by XGBoost, with no significant difference between them (Fig. 14b,c,e) when a larger dataset was compared to the data from an independent experiment (36.806e3; ∼1.22% of the full dataset) with extensive consideration of the spatial heterogeneity across the two benches. The major decrease in the accuracy of the Neural Network did not surprise us, as fluctuations until a steady state of accuracy could be reached were demonstrated previously (Fig. 11b). Here again, all of the algorithms were more accurate than the FPME. These findings substantiate our claim regarding the significance of spatial sampling in environmental research. Consequently, greater emphasis should be placed on the strategic distribution of measurement points, rather than simply increasing the quantity of sensors deployed or the volume of data amassed.

Our study demonstrates that machine-learning algorithms can predict evaporation more accurately than the FPME. This can be done using a relatively small dataset and small number of sensors, when the variability of the microclimates across the greenhouse is considered and the sensors are positioned accordingly. Future research should aim to determine whether microclimatic meteorological data and lysimeters can be used to predict plant transpiration, dynamic crop factors and yield. However, we expect that that task will be much more challenging due to the plant biological–environmental interactions of water-balance regulation mechanisms. We expect that the development of low-cost microsensors, utilizing wireless technologies, will provide better resolution of the dynamic plant–environment interaction, which will support valid measurements for machine-learning algorithms with higher degrees of accuracy for both spatial and temporal measurements.

## CRediT Author Statement

## Acknowledgements

The work was funded by the Israel Ministry of Innovation, Science and Technology (grant no. 0002122 and 001897).

## Supplementary

**Suppl. Fig. 1.**
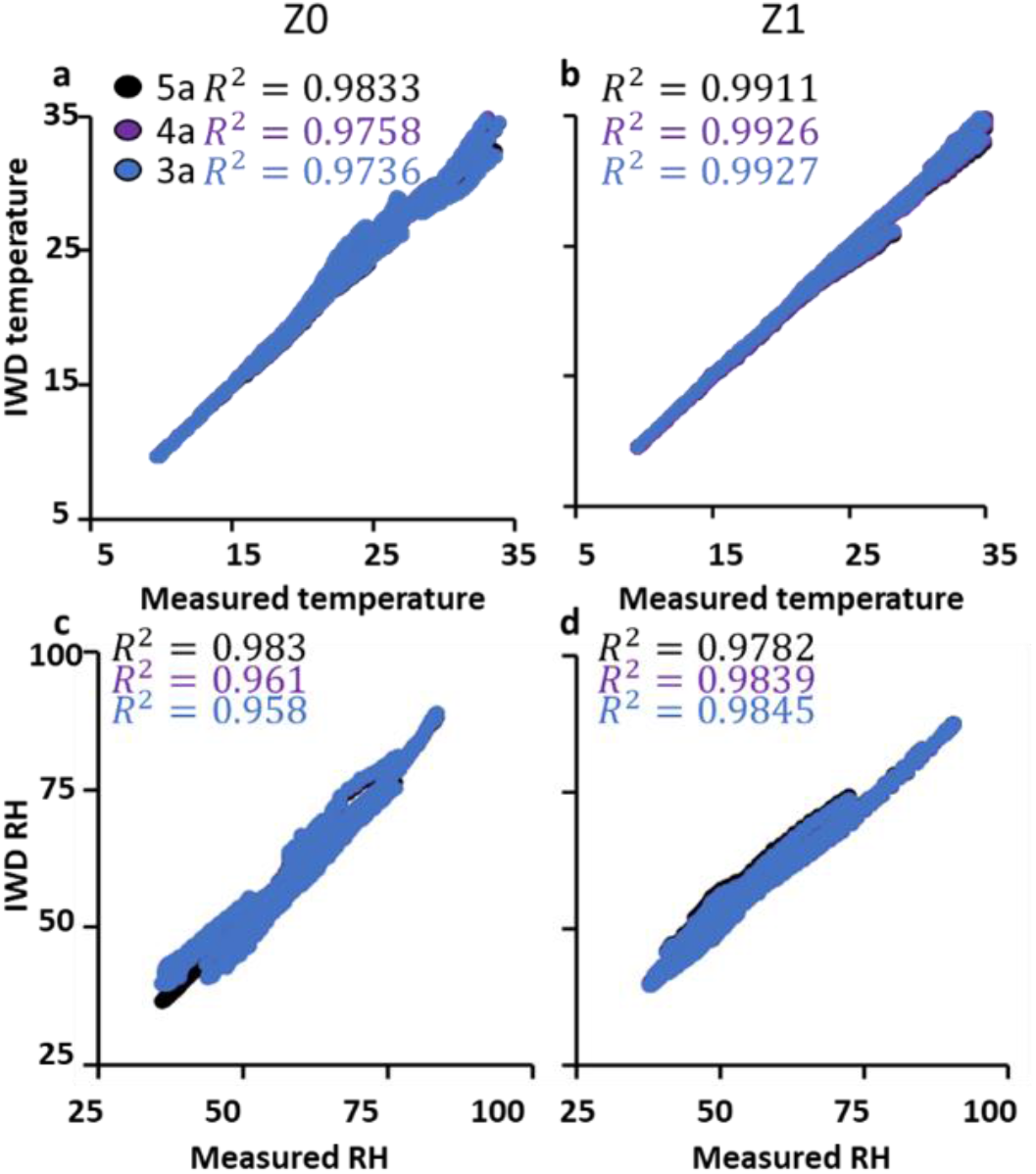
Correlations between calculated temperature and measured temperature, and between calculated relative humidity with measured relative humidity. We examined the correlation of measured meteorological data with IWD-calculated data for five, four and three MS units (5a, 4a and 3a, respectively) at a height of 0.3 m above the evaporating bath. All graph correlation *p*-values are <0.001 (tested with JMP).

**Suppl. Fig. 2.**
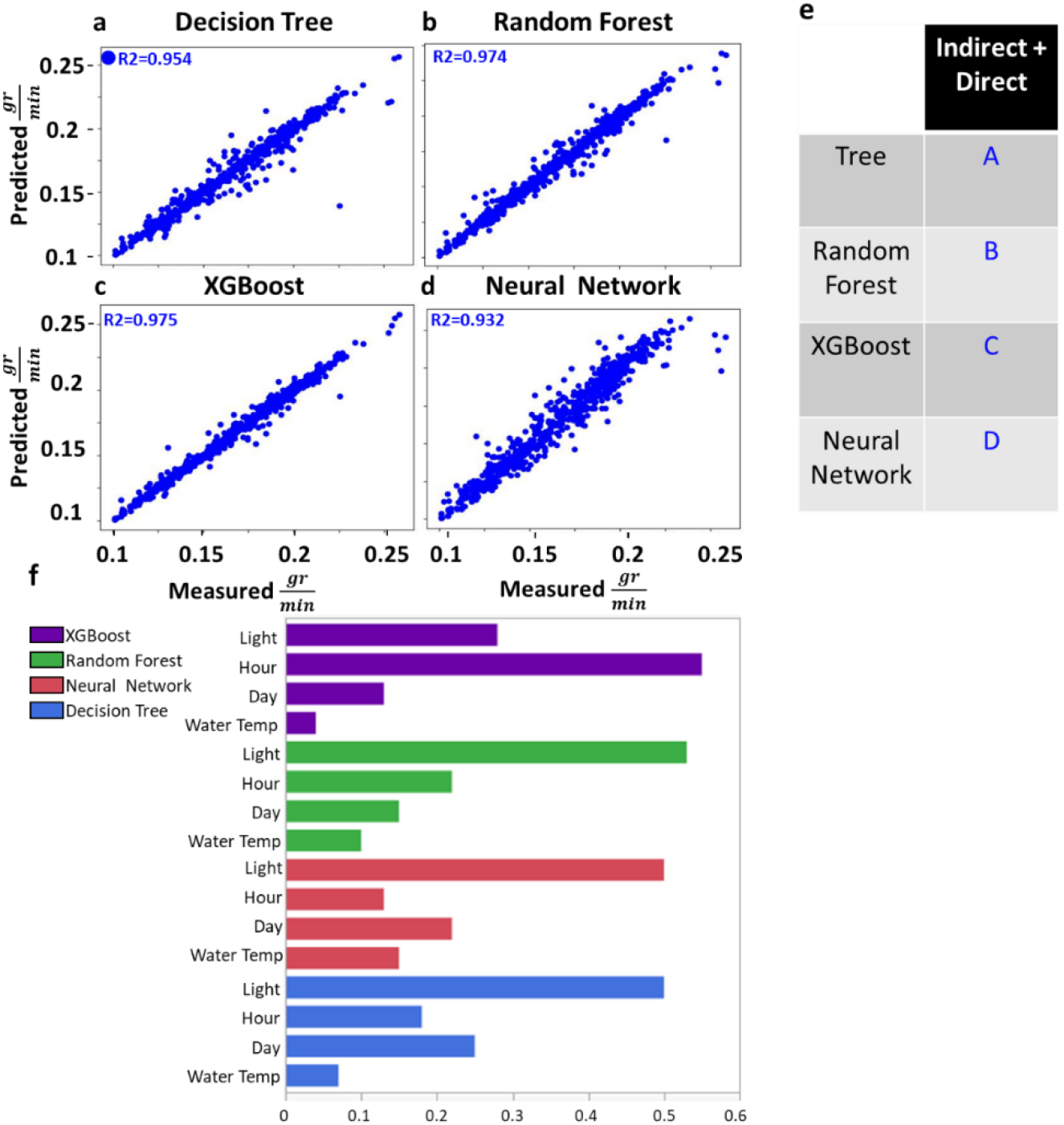
Comparison of algorithms based on water temperature, solar radiation and time data. A dataset including 1.2e6 data points for solar radiation, bath temperature and temporal parameters was used to evaluate the relationship between the measured evaporation rate and the predicted evaporation rate. All of the algorithms were applied to the same dataset: (a) Decision Tree, (b) Random Forest, (c) XGBoost and (d) Neural Network. (e) Connecting-letters report for the comparison of the errors observed for each of the algorithm models. (f) Feature hierarchy of the algorithms: Decision Tree (blue), Random Forest (green), XGBoost (purple) and Neural Network (red). (e) A connecting-letters report for the indirect and indirect + direct models, testing the difference of the model absolute value error between all of the predicted and the measured values, showed that the algorithms’ predictions were significantly accurate (*p* < 0.001). (f) Feature hierarchy of the machine-learning algorithms: Decision Tree (blue), Random Forest (green), XGBoost (purple) and Neural Network (red).

**Suppl. Fig. 3.**
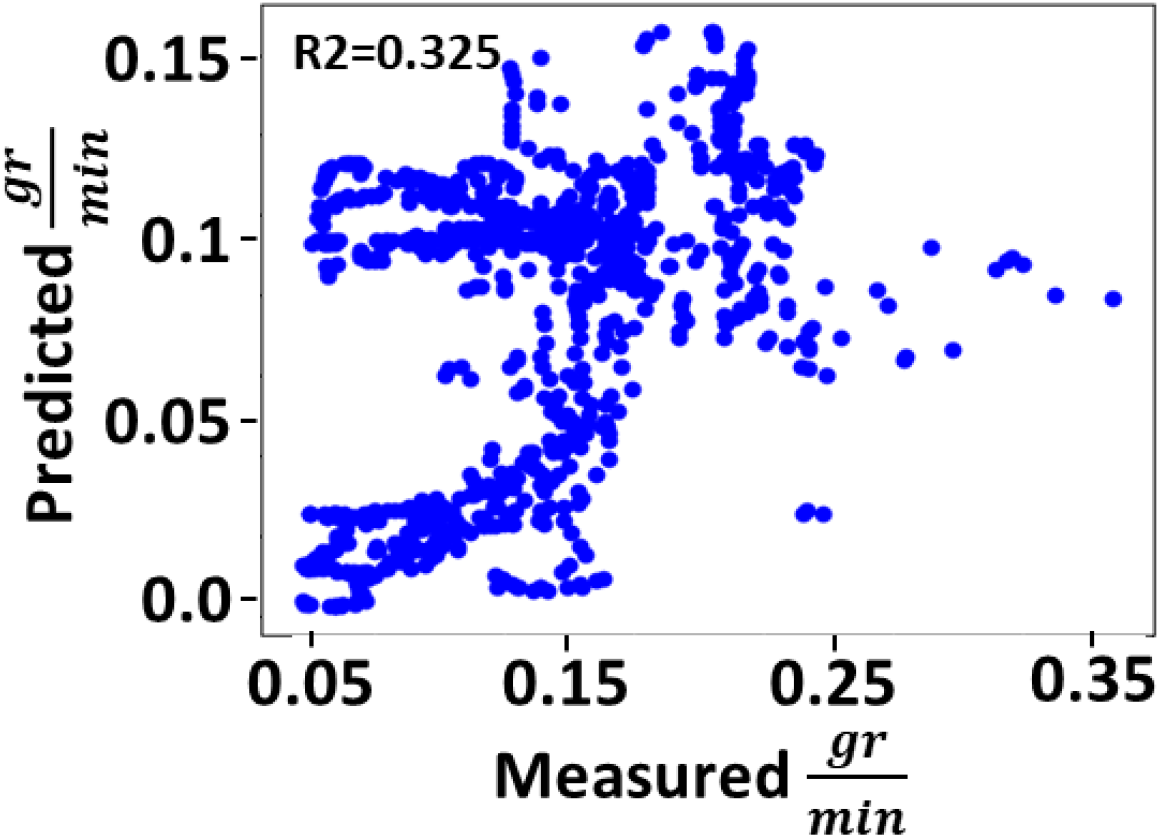
XGBoost prediction of the smaller dataset based on the larger-dataset model. A pre-trained machine-learning model based on the larger dataset was used to predict the result obtained using the smaller set of data (21.e3 data points) collected on the other side of the greenhouse. The correlation between the outputs obtained using the two datasets was not significant.

